# Revisiting Size Selective Mortality in Young Fish: Do Small Teleosts Really Pay a Cost?

**DOI:** 10.1101/2025.06.16.660013

**Authors:** Devon R. Bath, Levi Newediuk

## Abstract

Small juvenile and larval teleosts are typically more susceptible to starvation and predation, so the largest individuals tend to survive. However, most support for this ‘bigger is better’ hypothesis is experimental and does not account for the variation in food availability, competition, and other conditions that alter how starvation and predation affect small fish. To assess how natural populations experience size-selective mortality, we reviewed the past 30 years of literature on the subject while compiling 76 effect sizes of longitudinal survival data, which evaluates selection against size classes, to test for evidence that ‘bigger is better’. Our meta-analysis shows that the effect of body size on survival is consistently weak across species, populations, times, and locations. We discuss several reasons why larger young teleosts may not have a distinct survival advantage. We argue that they: 1) may be more profitable or noticeable to predators, 2) are unable to outgrow all their predators, and 3) require more resources, increasing their predator exposure out of necessity to forage. We recommend that mortality not be treated as constant across young teleost size classes, as different ecological conditions may favour smaller, larger, or neither size class. We suggest that the relationship between metabolic scope and mortality, geographic gradients in the importance of predation and starvation, and the interactive effects of alternative sources of mortality with predation and starvation in warming oceans are underexplored and could change how we think about the relationship between young teleost size and survival. Investigating these drivers will be important as temperatures rapidly rise, altering growth in wild fish.

## Introduction

Vulnerability to predation and starvation—the primary sources of mortality among larval and juvenile teleost fish—is influenced by an individual’s size (e.g., Garvey et al., 2004; Holmes & McCormick, 2010; van Deurs et al., 2011). Larger individuals are thought to face less predation risk because their size makes it harder for predators to capture and consume them (Juanes & Conover, 1994; Sogard, 1997). Larger fish are also more resistant to starvation as they tend to have more mass-specific stored energy and lower mass-specific metabolic rates (Auer et al., 2015). The idea that larger larval and juvenile fish are more likely to survive is the basis for the ‘bigger is better’ hypothesis. This hypothesis has been widely accepted in the literature for decades and forms the fundamental framework for understanding fish ecology (Bailey, 1984; Bailey & Batty, 1983; Bailey & Yen, 1983; Sogard, 1997).

While the ‘bigger is better’ hypothesis is influential in fish ecology, the relationship between size and survival is complex. Which prey sizes are most vulnerable to predation depends on which predators are present and their sizes relative to the prey (Juanes & Conover, 1994; Scharf et al., 1998). Behavioural dynamics among conspecifics can leave larger fish more vulnerable to predation and starvation (Polyakov et al., 2022; Reinhardt, 1999). Predators may target larger fish during shoaling because they are more noticeable (Polyakov et al., 2022). Importantly, ecological conditions can change seasonally and interannually, which previous studies note alter when size-selective mortality occurs and in what direction (Bath et al., 2024; Peres et al., 2022).

To understand the evidence for both sides of the size-selective mortality argument, we review evidence for and against the ‘bigger is better’ hypothesis, focusing on the two primary sources of mortality for young teleosts: starvation and predation (Gjøsæter et al., 2016; Sogard, 1997). First, we introduce the foundational ideas behind the hypothesis that led to widespread acceptance. We then synthesize longitudinal survivor data from studies of natural populations to test whether bigger fish enjoy lower rates of starvation and predation. Next, we review factors that influence the risk of predation and starvation, explaining why the ‘bigger is better’ hypothesis might hold in some cases and not in others. Our work builds on a foundational review by Sogard (1997), incorporating findings from recent studies to provide a contemporary consensus from the literature. Finally, we offer a perspective on areas of future research that would lead to our better understanding of the factors shaping early-life fish mortality and when the ‘bigger is better’ hypothesis applies.

### 1. Why is bigger better?

At the time of Sogard’s (1997) review of size-selective mortality in juvenile teleosts, evidence favoured the ‘bigger is better’ hypothesis. Here we discuss the ecological mechanisms for this pattern in larval and juvenile (hereafter, young) teleosts, which fall into two major categories: the metabolic efficiency of larger fish and their capacity for accessing and storing energy, which reduces their susceptibility to starvation, and the related behavioural and morphological adaptations that help them avoid predation.

Larger teleosts are more resilient to starvation than smaller ones because of physical advantages that help them store and access more energy. Endogenous energy storage scales with body size while mass-specific metabolic rate declines, allowing larger teleosts to store more energy while needing to forage relatively less (Biro et al., 2021; Brown et al., 2004; Shuter & Post, 1990). This means that during food shortages, larger teleosts can meet up to 80% of their metabolic needs from stored energy (Heintz & Vollenweider, 2010), leading to higher survival rates than smaller fish, approaching nearly 100% in some cases (Biro et al., 2021; Blom et al., 1994; Byström et al., 2006; Finstad et al., 2004; Geissinger et al., 2021; Pangle et al., 2005, 2011; Trexler et al., 1992). In contrast, smaller teleosts, with their high mass-specific metabolisms and limited energy reserves (Biro et al., 2021), lose body mass during long periods of low food availability because they lack sufficient endogenous stores (Fishelson, 1997; Heintz & Vollenweider, 2010; van Deurs et al., 2011).

Morphologically, body size improves foraging capacity in teleosts because gape limitations (i.e., how wide a fish can open its mouth) scale positively with size. Larger fish can consume a relatively wider range of prey sizes (Gaeta et al., 2018; Higgins et al., 2018; Pessanha & Araújo, 2014). Therefore, they are more likely to consume energetically profitable intermediate-size prey. Larger teleosts relative to others in the same development stages can also shift to more profitable prey guilds sooner, moving, for example, from invertebrates to other fish (Scharf et al., 2009). Improved diet quality during this transition reduces the need for teleosts to forage frequently, which lowers predation risk and increases energy gains, facilitating further growth. Additionally, the largest teleosts physically dominate others in their cohort and monopolize food sources, enhancing their growth rates and energy reserves (Beaugrand & Beaugrand, 1991; Naman et al., 2019; Reed et al., 2019; Whiteman & Côté, 2004).

The physiological and morphological advantages that help larger teleosts avoid starvation also translate into safer behavioural strategies that help them avoid predation. For example, lower mass-specific metabolic rates enable larger fish to reduce their foraging effort and avoid places where their predators are likely to hunt, choosing instead to remain in safer places with fewer resources (Garvey et al., 2004; Jonas & Wahl, 1998; Reinhardt & Healey, 1999; Sewall et al., 2019). In contrast, smaller teleosts store relatively less energy and use it less efficiently, forcing them to forage more often and in areas where predators are more likely to target them (Catano et al., 2016; Garvey et al., 2004; Heintz & Vollenweider, 2010; Naman et al., 2019; Pratt & Fox, 2002; Reinhardt & Healey, 1999; Sbragaglia et al., 2021). By physically dominating smaller teleosts, larger teleosts also control the resources in areas with fewer predators, further reducing their predation risk (Naman et al., 2019).

In addition to avoiding predation behaviourally, large teleosts are also harder to consume, making them less profitable prey. The largest teleosts can be inaccessible prey for other teleosts because they outgrow the maximum mouth width or ‘gape limit’ of predatory fish (Mihalitsis & Bellwood, 2017). Predatory fish and those from other guilds—including birds and invertebrates—also avoid eating larger prey due to longer handling times, which lower net energy gain and increase the risk of injury (Lucas et al., 2021; Lundvall et al., 1999; Mukherjee & Heithaus, 2013; Scharf et al., 1998; Taylor, 2003; Taylor & Collie, 2003; Tucker et al., 2016).

While there is good theoretical and experimental evidence for ‘bigger is better’, natural populations face ecological dynamics, such as fluctuating predation pressure and food availability, that can alter the benefits of size across time and space. Yet, studies in natural populations typically test size-selective mortality in only a few species, at ecologically similar sites, and over short timeframes (e.g., Pangle et al., 2011; Raventos & Macpherson, 2005; Takasuka et al., 2016). These studies also employ a variety of non-standardized methods that are difficult to compare directly (Sogard, 1997). As a result, we still lack a consensus understanding of how size affects survival in young teleost fish.

### 2. Meta-analysis of ‘bigger is better’

To improve our general understanding of size-selective mortality, we conducted a meta-analysis to assess size-selective mortality in survivor analysis studies, focusing on young teleosts. Survivor analysis, an established method to test for size-selective mortality in natural fish populations (Sogard, 1997), looks for relationships between survival and phenotypic traits. Longitudinal sampling compares the trait distributions between surviving individuals and an initial sample of young teleosts (Figure 1). If survivors differ phenotypically from the initial sample of young teleosts, we assume that selection for that trait led to non-random mortality. In size-selective mortality studies, size is the focal trait in the survivor analysis. A single group of young teleosts is measured at two points in time, and support for ‘bigger is better’ is concluded if the average length of surviving individuals is greater than the original sample (Figure 1).

**Figure 1.**
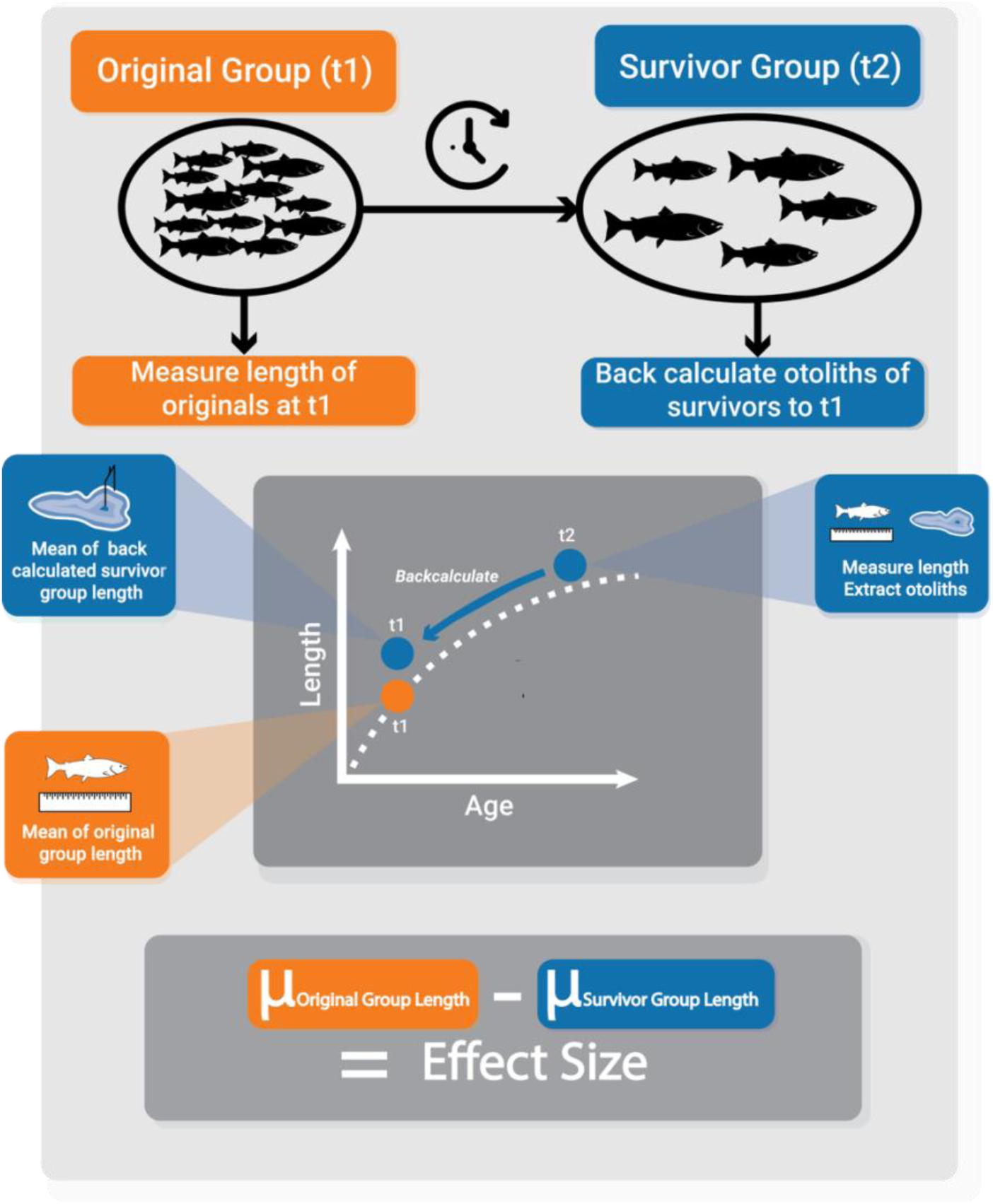
Schematic of effect size calculations derived from survival analysis. The original group represents the first capture at time 1 (t1), and the survivor group represents the individuals that survived to a later capture at time 2 (t2). The length of individuals in the original group and their mean length at t1 are measured. Mean and individual lengths of the survivor group at t2 are derived from extracted otoliths using the known relationship between otolith and somatic growth. The effect size is calculated by subtracting the mean lengths of the original group (µ_survivor group length_) from the survivor group (µ_original group length_).

## Methods

In our literature search, we used the search terms ‘size selective mortality’ and ‘bigger is better’, as these terms are common keywords in survival analysis studies. First, we systematically searched the Web of Science core collection on February 12th, 2024. Then, we searched for papers on the Dryad data repository on April 15th, 2024, employing an application programming interface in Python. Search details are in S1 Methods.

We selected studies based on several requirements. First, we included studies focused on our two target life history stages of young teleost—larval or juvenile. To facilitate our effect size calculations—i.e., the direction and magnitude of effect standardized across studies—we only included studies that measured the mean length of an initially sampled cohort of fish, i.e., the ‘original,’ and the back-calculated mean length of a surviving group of fish using the otoliths (Figure 1). Additionally, we only included studies that reported sample size for the original and survivor groups and a measure of variation for each group. We placed no constraints on the years that papers were published.

To calculate the effect size across studies, we used ‘C*ohen’s d’*, i.e., the standardized mean difference between the survivor and original groups (Harrer et al., 2021). We calculated effect sizes so that negative values indicated selection against smaller young teleosts and positive values indicated selection against larger individuals. When studies compared multiple survivor groups with a single original group, we considered them separate effect sizes. We considered comparisons in different years, locations, and from different species as separate effect sizes, even when they came from the same studies.

Our initial search returned 829 publications. After screening the publications by journal, title, and abstract (S1 Methods), we found 10 studies were suitable for our meta-analysis (Figure 2). From these studies, we could extract 76 effect sizes representing 16,264 individual fish. The effect sizes represent 10 species from 6 orders of teleosts, including Salmoniformes, Blenniiformes, Clupeiformes, Labriformes, Perciformes and Gobiiformes (Table S3). The phylogeny covers species that are anadromous, freshwater, and marine. Larval teleosts represent 79% of the effect sizes and juvenile life stages, 21% (Table S4). The effect sizes are from the northern hemisphere only, but their geographic locations span broadly east and west of the prime meridian (Figure 3). The studies also represent several life events, including marine migration, metamorphosis, overwintering, post-hatching, post-settlement, and young of the year (Table S3).

**Figure 2.**
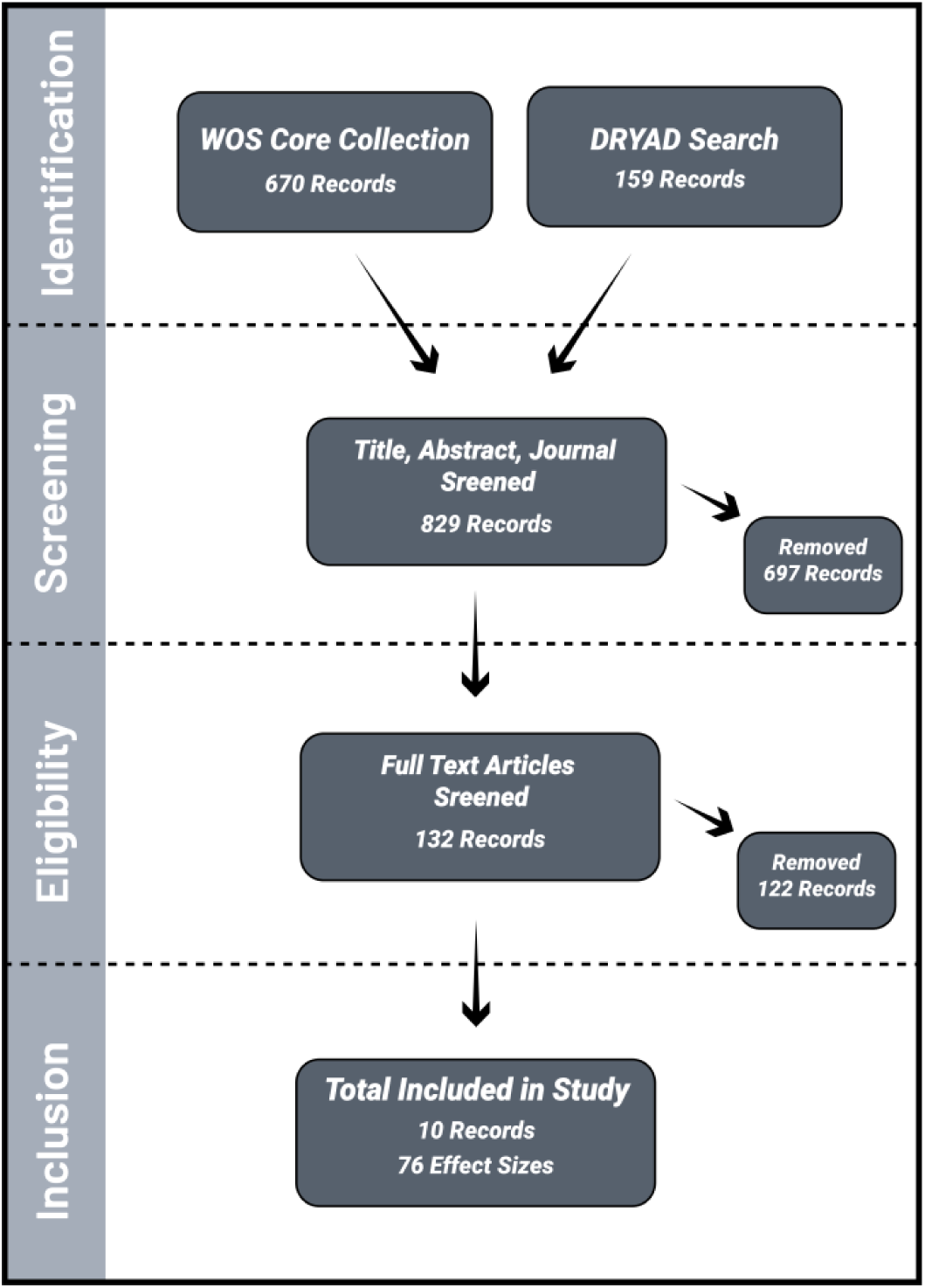
PRISMA diagram outlining our criteria for study inclusion in the meta-analysis. In the identification stage, we searched the Web of Science Core Collection and DRYAD data repository for studies matching the keyword terms ‘size selective mortality’ and ‘bigger is better’. We screened all publications at the abstract stage based on life history stage, method, and required data (‘screening’), and eliminated all studies not matching our criteria after a full-text reading (‘eligibility’). We included studies from which we could calculate at least one effect size (‘inclusion’).

**Figure 3.**
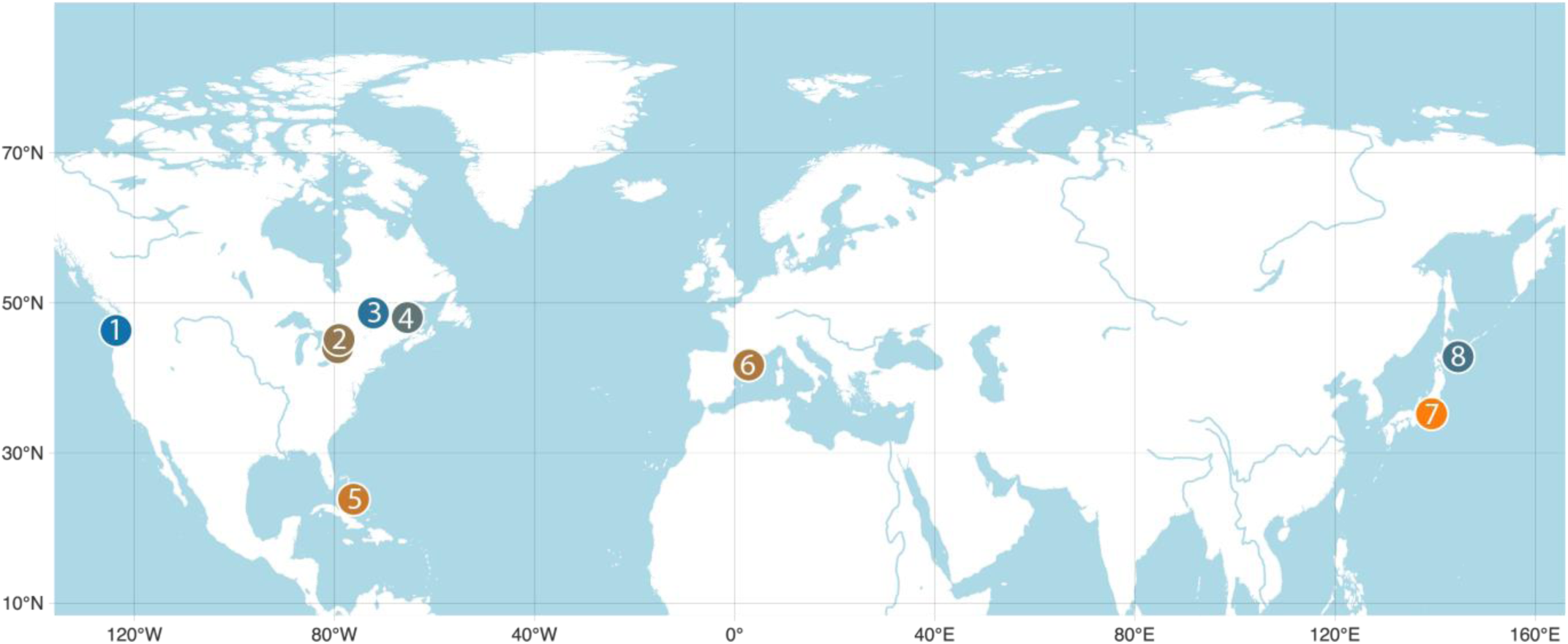
Map showing the sampling locations of studies included in the meta-analysis. Each coloured point is a different study, several of which represent multiple effect sizes. The map is displayed using unprojected geographic coordinates (WGS 84, EPSG:4326), with longitude and latitude in decimal degrees. 1. Claiborne et al., 2011; 2. Post & Prankevicius, 1987; 3. Good et al., 2001; 4. Johnston et al., 2005; 5. Samhouri et al., 2009; 6. Raventos & Macpherson, 2005; 6. Macpherson & Raventos, 2005; 7. Takasuka et al., 2004; 7. Takasuka et al., 2016 ; 8. Honda et al., 2020.

To estimate true effect sizes for studies and the pooled effect between studies, we fit a Bayesian, intercept-only mixed-effects model using the package ‘brms’ (Bürkner, 2017) in R (R Core Team 2023), specifying study ID as the random effect. We used an intercept-only model because we are only interested in the relationship between body size and survival.

### Results

Our meta-analysis, including 16,264 individuals across 10 studies, showed a weak effect of size on survival. The pooled effect size across studies was -0.056 (95% CI: -0.14 to 0.035; Figure 4). The probability that the pooled effect size is less than 0 was 90% (Figure S1). This means that the pooled effect size has a 90% probability of being negative, weakly supporting greater survival for larger fish. Except for one study, the credible intervals of all effect sizes crossed zero, which indicates a similarly weak relationship between body size and survival in young teleost fish regardless of year, location, or species.

**Figure 4.**
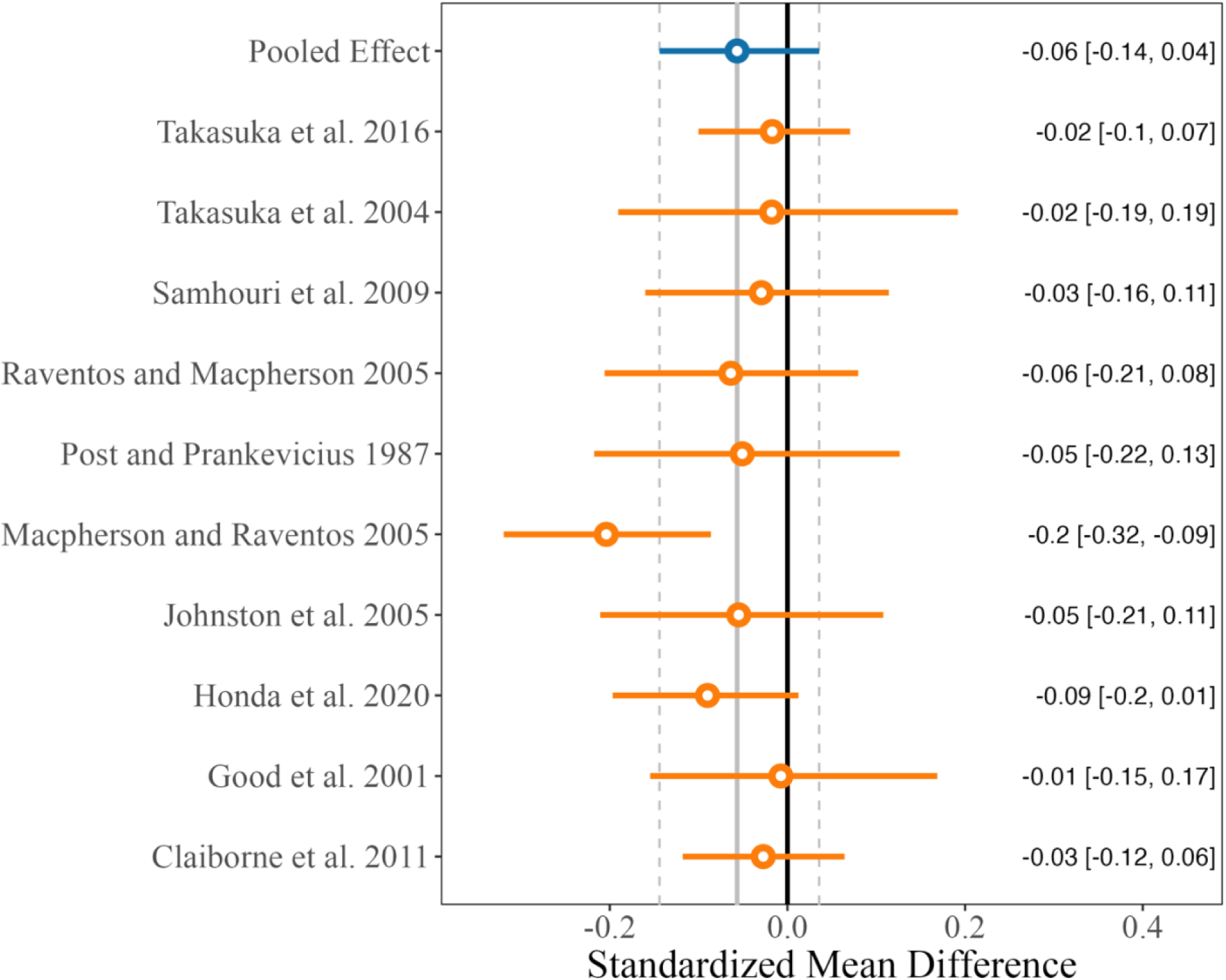
Forest plot of study true effect sizes and the pooled effect size of survivor analyses where a negative effect size indicates larger individuals had greater survival. Length of lines denote the upper and lower 95% credible interval. The effect size is annotated on the right-hand side with the 95% credible interval in square brackets. The grey solid line represents the true pooled effect size, and the dashed grey lines represent the 95% credible interval of the pooled effect size.

A single study, Macpherson & Raventos (2005), accounted for 47% of all effect sizes in our meta-analysis. The effect size from this study was the largest (−0.20), with the only confidence interval not crossing zero (95% CI: -0.31 to -0.077; Figure 4; Table S2). To ensure our results were not biased by this large study, which included multiple species, years, and locations (S1 Methods), we excluded the study and re-ran the model. Its removal increased the pooled effect size from -0.056 (95% CI: -0.14 to 0.035) to –0.022 (95% CI: -0.61 to 0.027; Figure 5), and the probability of the pooled effect size being less than 0 decreased to 76%, but the pooled effect size did not change direction. This means the weak relationship we found between body size and survival persisted across the other less influential studies.

**Figure 5.**
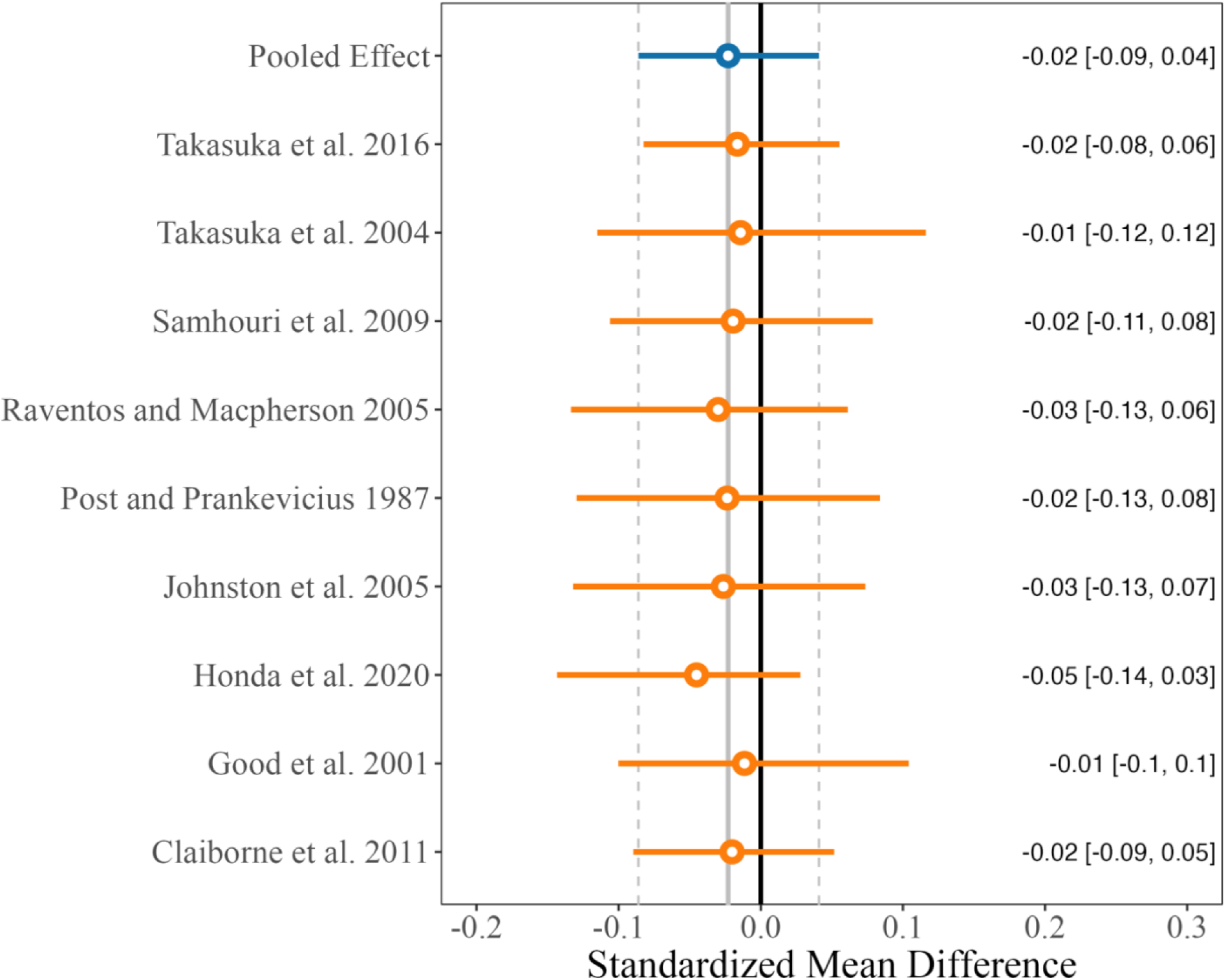
Forest plot of the sensitivity analysis containing study true effect sizes and the pooled effect size of survivor analyses with the largest study removed. Negative effect sizes indicate larger individuals had greater survival. Length of lines denote the upper and lower 95% credible interval. The effect size is annotated on the right-hand side with the 95% credible interval in square brackets. The solid grey represents the pooled effect size and the dashed grey lines represent the pooled effect size’s 95% credible interval.

Despite a consistently weak relationship between size and survival in our meta-analysis, given the large spread in true effect sizes across studies, our results also suggest that the exact strength of selection against smaller fish still depends on the time, location, and species being tested (Table S1). This finding supports similar observations from Sogard (1997), indicating that bigger is not always better, and hinting that factors other than size could help explain the additional variation in survival among young teleosts.

### 3. Why is bigger not always better?

Although many studies report that larger fish experience higher survival rates, this is not always the case in natural populations because predator-prey relationships are complex. Under optimal foraging, fish predators should select prey sizes that maximize net energy gain (Charnov, 1976; Sogard, 1997; Werner & Hall, 1974; Werner & Mittelbach, 1981). However, which prey sizes maximize energy gain also depend on predator traits. When young teleosts face diverse predators under natural conditions, predation pressure should be distributed more evenly across size classes, leading to no clear size advantage (Figure 6A). Additionally, young teleosts, regardless of size, are willing to risk predation to access the resources they need when growing (Catano et al., 2016; Naman et al., 2019; Sbragaglia et al., 2021).

**Figure 6.**
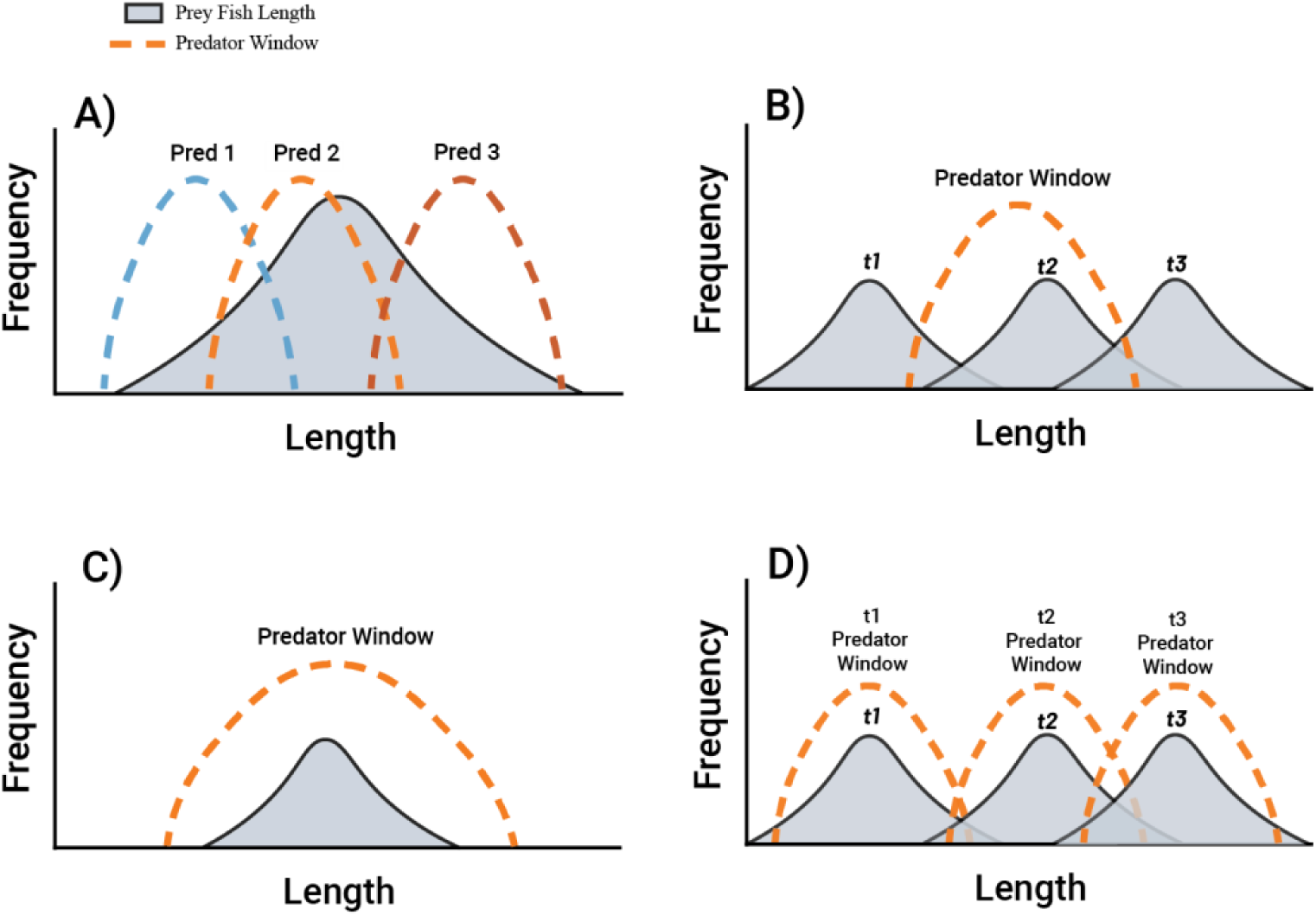
Conceptual figure showing the relationship between young teleost size and survival under different predation contexts. Orange, blue, and red dotted lines represent a predator’s consumption window for a particular prey size frequency distribution (grey). **A)** Shows how the presence of multiple predators could cause stabilizing selection by selecting prey across the size frequency distribution. **B)** Indicates how a fish growing over time (t1 to t3) could move through a predator window, changing the direction of size-selective mortality over time. **C)** Displays how a predator could encompass an entire size frequency distribution of prey, and **D)** demonstrates how a predator could synchronize growth rates with a prey, in both cases eliminating size-selective mortality.

Prey vulnerability to predation shifts over time and across locations based on predator traits. Predator capture success and preferences often follow a dome-shaped distribution of prey sizes that depends on the predator’s size (Cowan et al., 1996; Sigler et al., 2009). Thus, size-selective morality is not dependent on the size of the young teleost relative to others but on predator size preferences (i.e., optimal foraging; Charnov, 1976; Scharf et al., 1998, 2009; Sigler et al., 2009; Taylor, 2003). This means prey species might face selection against smaller or larger individuals or no size-selective mortality at all, depending on which predators are present (e.g., Holmes & McCormick, 2006, 2010; Figure 6A). As young teleosts grow, their value to predators also changes, so selection patterns can change over time (Figure 6B). For example, predators removed the largest damselfish (*Pomacentrus amboinensis*) on their initial settlement, but predation shifted to target the smallest individuals one month later (Meekan et al., 2010).

Young teleosts face multiple predators, each of which influences the advantages of different sizes. For larger individuals to consistently benefit from reduced predation risk, most predators would need to prefer smaller prey. However, under natural conditions, co-occurring predators often specialize on different prey sizes (Holmes & McCormick, 2010; Figure 6A). When predators are removed experimentally, the survival advantage shifts toward prey sizes favoured by the removed predator (McCormick & Hoey, 2004), highlighting that the survival advantage of large young teleosts is highly dependent on the predator pool.

Young teleost fish can outgrow their predators, but they may be unable to do so when the predator’s preferred size window encompasses the entire size distribution of its prey (Robert et al., 2010; Figure 6C). Some predators have adapted to grow synchronously with the prey species they consume, ensuring their prey remains within an optimal size range for consumption (e.g., Persson & Bronmark, 2002; Figure 6D).

Some stages of development are energetically expensive, changing the relationship between size and foraging risk. While energy stores usually scale with size, at earlier life stages, fish devote more energy to growth, reducing energy storage, and resulting in larger young teleost fish with lower energy stores (Abookire et al., 2024; Biro et al., 2005; Schultz & Conover, 1997; Sewall et al., 2019). Transitioning from the larval to juvenile stage can be energetically expensive, requiring as much as half of a fish’s energy reserves (Nursall & Turner, 1985; Youson, 1988). These developmental changes mean that young teleosts in the early juvenile stage can often have lower lipid levels than smaller individuals that are in the late larval stage. Individuals with few energy stores are at risk of starvation, and as a result, they are often more willing to risk predation while foraging (Dall et al., 2004). As a result of this risk-taking behaviour, predators may opportunistically select larger prey instead of smaller ones (McCormick & Hoey, 2004).

Some of the collective strategies young teleosts use to lower their predation risk may increase risk to the largest individuals in the group. Shoaling behaviour, for example, lowers individual risk if all individuals in the group are similar in size. However, individuals larger than the average shoal member attract predators, increasing their mortality rates (Polyakov et al., 2022). Also, individual behavioural traits like boldness, which determine risk-taking in fish, often increase with size (Brown et al., 2007). This means larger young teleosts might be more willing to accept risk while foraging, resulting in greater predation of larger individuals (Biro et al., 2005; Stamps, 2007).

While larger young teleosts typically dominate access to resources, which helps them survive resource limitations, they may be more vulnerable to starvation when under predation risk. Larger fish use dominance behaviours to monopolize resources, as their greater standard metabolic rates (Metcalfe et al., 2015; Reid et al., 2012; Yamamoto et al., 1998) require them to consume more energy. However, the presence of predators suppresses these dominance behaviours in larger fish, reducing their food intake relative to their mass and increasing their risk of starvation (Cutts et al., 2002; Johnsson, 1993; Metcalfe et al., 2015; Naman et al., 2019). Also, suppressing dominance behaviours raises predation risk to larger individuals, as they may resort to riskier foraging behaviours than smaller individuals to meet their energy needs for growth (Johnsson, 1993; Metcalfe et al., 2015).

Although previous experiments and field work have suggested larger young teleosts have higher survival rates, the relationship between body size and survival is less clear under natural conditions. The vulnerability of different size classes to predation varies depending on the predator size and the composition of the predator community. Additionally, larger young teleosts remain susceptible to starvation, as energy storage is not always proportional to size, and dominance hierarchies break down under high predation pressure. These dynamics in natural predator-prey communities likely explain the absence of a strong size-selective mortality signal we observed in our meta-analysis.

## Literature gaps

In the previous section, we outlined several ecological contexts in which predation patterns in young teleost fish might not be size selective. Predicting when these ecological contexts might arise will require uncovering the relationship between size, metabolism, and dominance in fish, mapping geographic patterns of size-selective mortality in response to the environment, and considering the additive effects of mortality sources other than starvation and predation. In this section, we describe three future directions for research in these key areas.

Studying them should improve our overall knowledge of fish ecology and population dynamics, particularly as the climate warms.

### Gap 1—The link between metabolic scope and size-selective mortality

Metabolic scope, which typically increases with size, is thought to influence size-selective mortality in young teleosts (Clarke & Johnston, 1999; Jerde et al., 2019). This is because metabolic scope can determine access to resources, vulnerability to predation, and ultimately survival (Killen et al., 2006; Naya & Bozinovic, 2012). However, it is unclear whether these factors are actually determined by metabolic scope or resting and maximum metabolic rates. Because metabolic scope is the difference between resting and maximum metabolic rates, individuals with the same metabolic scopes may still differ greatly in their metabolic profiles (Norin & Malte, 2012).

Resting and maximum metabolic rates may be the key drivers of size-selective mortality for several reasons. First, having a higher resting metabolic rate can mean a greater energy demand, which is often considered the driver of dominance and risk-taking behaviours in young teleosts that affect mortality (e.g., Auer et al., 2015; Cutts et al., 2002). Furthermore, regardless of metabolic scope, maximum metabolic rate can restrict the behaviours young teleosts use to acquire energy. Thus, to understand how size affects survival, we need to know the extent to which behavioural differences related to specific metabolic profiles drive mortality in young teleosts. These studies will be best carried out under natural conditions where predators are present (Reinhardt, 1999) and food patches and habitats are complex (Reid et al., 2012), providing a realistic picture of how metabolic scope relates to behaviour and size-selective mortality.

### Gap 2—Geographic gradients

The relationship between size and survival likely operates on a latitudinal gradient. At higher latitudes, survival in young teleosts is influenced by shorter growing seasons and periods of low food availability (Shuter & Post, 1990). Their size and lipid amount often increase with latitude (Knouft, 2004; Saunders & Tarling, 2018; Schultz & Conover, 1997), as reaching a size with adequate energy stores can reduce the risk of starvation and minimize foraging-related predation (Garvey et al., 2004; Jonas & Wahl, 1998; Sewall et al., 2019; van Deurs et al., 2011). This means that size-selective mortality due to starvation might be more important for young teleosts at higher latitudes. Conversely, young teleosts at lower latitudes should be less prone to starvation due to higher resource availability but may face greater predation risk from a more diverse predator pool. When multiple predators are present in a diverse predator pool, predator selection across the size-frequency distribution of prey could cause stabilizing selection on size, where greater size does not increase survival.

Distinguishing the most influential source of mortality—predation or starvation—is crucial for understanding how and when size should influence population dynamics. While some models assume constant mortality across size classes (e.g., Froese et al., 2008), this assumption may not hold across environmental gradients. Testing for size-selective mortality across latitudes, where predator communities and resource availability vary, could reveal whether this assumption is consistent. This is especially important since climate change will likely continue to alter size-selective mortality in young teleosts.

### Gap 3—Additive and interactive effects

Our review focused on two sources of mortality in young teleosts: predation and starvation. However, other equally important sources can compound mortality risk, sometimes with additive effects for larger individuals. For example, parasites often affect smaller fish disproportionately (e.g., Smith et al., 2022; Tucker et al., 2002). However, larger individuals can develop higher parasite loads over time because of their larger surface areas for infection, and underdeveloped immunocompetence, which they trade off for faster growth rates (Bartuseviciute et al., 2022). Additionally, temperature can compound foraging risk for smaller fish by elevating their metabolic demand (Pink & Abrahams, 2016) and increasing predator activity.

The future impact of climate warming on size-selective mortality through interactive and additive factors will be a key area of future study. The transmission and biodiversity of aquatic parasites in warmer waters are highly uncertain (Lõhmus & Björklund, 2015). Also, the shrinkage of body size in fish caused by global temperature increases will have important and complex effects on size-selective mortality (Daufresne et al., 2009; Deutsch et al., 2022; Forster et al., 2012). For example, while temperature could shift size-selective mortality distributions, fish predators are also reaching smaller sizes under higher global temperatures, which will relieve some predation pressure (Eskuche-Keith et al., 2024). Thus, exploring the mechanisms by which additive and interactive sources of mortality, like parasites and temperature, interact with size-selective mortality from predation and starvation, in both experimental and natural settings, will help anticipate and explain unexpected relationships between size and mortality in young teleosts.

## Conclusion

We provide evidence that, as a young teleost, being bigger may hold only a weak advantage in the face of mortality. Assumptions of the ‘bigger is better’ hypothesis, while well supported experimentally, are limited by complex interactions under natural conditions. In environments where predators prefer different-sized prey, all sizes of young teleost may be vulnerable. However, as climate change alters community composition and maximum growth, the relationship between size and vulnerability is likely to change. We suggest that future longitudinal studies under natural conditions consider how latitude, metabolic physiology, and additive effects might change the relationship between size and mortality for young teleosts. This understanding will help anticipate mortality patterns under new conditions caused by the changing climate.

