## Supplementary for "Revisiting Size Selective Mortality in Young Fish: Do Small Teleosts Really Pay a Cost?"

### S1. Methods

*Intercept only model and prior distributions*

We created a Bayesian intercept-only mixed-effects model using the package 'brms' (Burkner et al. 2021) in R (R Core Team 2023) to estimate a true effect size across studies. Studies were used as random intercepts for the model. Intercept-only models estimate the pooled true effect size, the study true effect size, variation within study true effect size and variation between study true effect sizes (i.e., between study heterogeneity).

We used regularizing priors to estimate the mean of true effect sizes, *~N(0,1)*. A normal prior distribution with a mean of 0 and a standard deviation of 1 specifies a 95% prior probability that the true pooled effect size lies between –2.0 and 2.0. We assigned a Half-Cauchy prior for between-study heterogeneity with location parameter = 0 and scaling parameter = 0.5, *~Half-Cauchy(0, 0.5)* (Harrer et al. 2021). We calculated effect sizes using the standardized mean difference between the mean of the original sample and surviving fish lengths, i.e., *Cohen's d*, (see Figure 1 in the main text for a schematic). Specifying a half-Cauchy prior distribution with a scaling parameter of 0.5 ensures the between-study heterogeneity can take on values near 0 while allowing an estimation of values that may be considered higher than typical for a meta-analysis (Harrer et al. 2021).

*Sensitivity analysis*

To ensure our meta-analysis results were robust to the disproportionate influence of the largest study, we also conducted a sensitivity analysis where we removed the largest study, Macpherson and Raventos (2005), with 32 of 76 effect sizes accounting for 42% of all effect sizes in our meta-analysis. Macpherson and Raventos (2005) conducted survivor analysis of two species of reef fish (*Lipophrys trigloides* and *Chromis chromis*). Survivors were captured at the juvenile stage and back-calculated to both time at larval settlement and size at hatching. Measurements were made at four time points over two years, resulting in 32 effect sizes in total.

*Model convergence and fit*

We inspected all model trace plots for discernible trends that could indicate non-convergence. In addition, we checked that the scale reduction factor, i.e., the Gelman-Rubin diagnostic, was less than 1 (Brooks and Gelman 1998). Finally, we performed posterior predictive checks that the fitted model error distributions matched the error distributions of the observed data.

*Between-study heterogeneity and publication bias*

Between-study heterogeneity is the variation within the pooled true effect size due to differences between study true effect sizes. We directly estimated the between-study heterogeneity using the Half-Cauchy distribution.

We tested for publication bias by visually inspecting a funnel plot of the pooled standard error of the observed effect sizes. Larger effect sizes than expected for the standard error, which contribute to a lop-sided funnel shape, indicate that positive results may be published more often than null results.

### S2. Results

*Study Search*

We screened 829 articles, including 670 from our initial Web of Science (WOS) search and 159 from the Dryad Data Repository (hereafter Dryad). We initially identified unsuitable WOS studies based on publication titles indicating work outside the field, e.g., the Irish Veterinary Journal, International Journal of Cardiology, etc. We used the function '*grepl'* in R (R Core Team 2023) to remove 248 irrelevant articles by journal title, leaving 581 articles for manual screening. We next screened article titles, removing 287 articles for having titles we identified as unsuitable, leaving 294 potentially suitable articles. We screened the remaining articles by abstract, removing 162 articles, leaving 132 for comprehensive screening. None of the three potentially suitable articles from the Dryad search were usable, and 10 from the WOS were considered usable. We contacted the authors of 35 additional potentially suitable articles for raw data or additional information needed to estimate effect sizes. Unfortunately, all such attempts were unsuccessful.

*Intercept only model*

There was weak evidence of greater survival in larger juvenile teleosts. The pooled effect size across studies is -0.056 (95% Credible Interval: -0.14 to 0.035; Table 1; Figure 1). The between-study heterogeneity (0.09; 95% CI: 0.03 to 0.18; Figure 2) was nearly double the pooled effect size (-0.056), indicating that the true effect size across studies varied greatly (Table S1). Within study heterogeneity (0.01; CI: 0.00 to 0.04; Figure S3) was nearly one-fifth of the pooled effect size, indicating little variation within studies (Table 1). The overall effect of size-selective mortality on young teleosts appeared weak, but the high between-study heterogeneity indicates that ecological factors specific to a study area likely led to variation in size-selective mortality among sites.

*Sensitivity analysis*

The pooled effect size estimated in the sensitivity analysis (i.e., the model with the largest study removed) was larger than the original model (–0.022; 95% CI: -0.61 to 0.027; Table S2; Figure S4). Between-study heterogeneity was lower in the sensitivity analysis (0.03; 95% CI: 0.00 to 0.09; Figure 5), and the pooled effect size was approximately double that of the original model, indicating substantial variation in the true effect size of studies (Table S2). Within study heterogeneity was approximately the same as the original model (0.01; CI: 0.00 to 0.04) but was nearly three-quarters of the pooled effect size, indicating moderate variation within studies (Table S2; Figure 6).

*Publication Bias, Model Convergence and fit*

Asymmetry in the funnel plot of observed effect sizes followed a negative skew (Figure S7), indicating possible bias in publishing results that support the 'bigger is better' hypothesis. Both the original model and the sensitivity analysis model converged. R-hat (scale reduction) values were 1 for both models, and trace plots indicated that MCMC chains adequately explored the parameter space (Figure S8). The posterior predictive checks indicated our model fit the observed data well (Figure S9).

#### Tables

**Table S1.** Intercept-only model estimates, estimate error, and 95% credible intervals (CI) are shown for the pooled effect size, between-study heterogeneity, and within-study heterogeneity of the original model with all studies retained.

| | Estimate | Error | CI 2.5% | CI 97.5% |
|---|---|---|---|---|
| Pooled effect size | -0.057 | 0.04 | -0.12 | 0.029 |
| Between study heterogeneity | 0.098 | 0.04 | 0.03 | 0.20 |
| Within study heterogeneity | 0.027 | 0.01 | 0.00 | 0.08 |

**Table S2.** Intercept-only model estimates, estimate error, and 95% credible intervals (CI) are shown for the pooled effect size, between-study heterogeneity, and within-study heterogeneity of the sensitivity analysis model with the largest study removed.

| | Estimate | Error | CI 2.5% | CI 97.5% |
|---|---|---|---|---|
| Pooled effect size | -0.022 | 0.02 | -0.06 | 0.03 |
| Between study heterogeneity | 0.04 | 0.03 | 0.00 | 0.13 |
| Within study heterogeneity | 0.03 | 0.02 | 0.00 | 0.07 |

**Table S3.** Information from studies included in the meta-analysis. Studies are characterized according to their name, the species that are tested, the taxonomic order, the life history stage when measured, what life event occurred while testing according to the study authors, migration pattern, latitude, and longitude of sampling, the number of effect sizes of a given row and the pooled sample size. The pooled sample size refers to the sample size of the initial or original group sampled and the sampled surviving group.

| ***Study name*** | ***Species*** | ***Order*** | ***Life history stage*** | ***Life event*** | ***Migration pattern*** | ***Latitude*** | ***Longitude*** | ***Number of effect sizes*** | ***Pooled sample size*** |
|---|---|---|---|---|---|---|---|---|---|
| Claiborne et al. 2011 | Oncorhynchus tshawytscha | Salmoniformes | Juvenile | Marine migration | Anadromous | 46.3 | 123.7 | 4 | 9655 |
| Good et al. 2001 | Salmo salar | Salmoniformes | Juvenile | Post hatching | Anadromous | 48.3 | 69.9 | 2 | 316 |
| Honda et al. 2020 | Oncorhynchus keta | Salmoniformes | Juvenile | Marine migration | Anadromous | 42.8 | 144.6 | 6 | 1366 |
| Johnston et al. 2005 | Salmo salar | Salmoniformes | Juvenile | Overwinter | Anadromous | 48 | 65.5 | 1 | 75 |
| | Salmo salar | Salmoniformes | Larvae | Overwinter | Anadromous | 48 | 65.5 | 1 | 79 |
| | Salmo salar | Salmoniformes | Juvenile | Overwinter | Anadromous | 48.2 | 65.8 | 1 | 55 |
| | Salmo salar | Salmoniformes | Larvae | Overwinter | Anadromous | 48.2 | 65.8 | 1 | 60 |
| Macpherson and Raventos 2005 | Chromis chromis | Blenniiformes | Larvae | Post settlement | Marine | 41.7 | 2.8 | 20 | 816 |
| | Lipophrys trigloides | Blenniiformes | Larvae | Post settlement | Marine | 41.7 | 2.8 | 16 | 644 |

| | | | | | | | | | |
|---|---|---|---|---|---|---|---|---|---|
| Post and Prankevicius 1987 | Perca flavescens | Perciformes | Juvenile | Young of the year | Freshwater | 44 | 79.4 | 1 | 143 |
| | Perca flavescens | Perciformes | Juvenile | Young of the year | Freshwater | 45.1 | 79.1 | 1 | 129 |
| Raventos and Macpherson 2005 | Symphodus ocellatus | Labriformes | Larvae | Post settlement | Marine | 41.7 | 2.8 | 2 | 198 |
| | Symphodus roissali | Labriformes | Larvae | Post settlement | Marine | 41.7 | 2.8 | 2 | 204 |
| Samhouri et al. 2009 | Gnatholepis thompsoni | Gobiiformes | Larvae | Post settlement | Marine | 23.8 | 76.2 | 8 | 515 |
| Takasuka et al. 2004 | Engraulis japonicus | Clupeiformes | Larvae | Metamorphosis | Marine | 35.2 | 139.3 | 1 | 131 |
| Takasuka et al. 2016 | Engraulis japonicus | Clupeiformes | Larvae | Metamorphosis | Marine | 35.2 | 139.3 | 9 | 1878 |

**Table S4.** Pooled sample sizes and number of effect sizes (n) summarized by category: study name, life event, migration pattern, life history stage, and species. Percentages indicate the percentage of effect sizes and pooled sample sizes from each category.

| **Study name** | | **Pooled sample size** | **Number of effect sizes** |
|---|---|---|---|
| | Claiborne et al. 2011 | n = 9,655 (59.4%) | n = 4 (5.3%) |
| | Good et al. 2001 | n = 316 (1.9%) | n = 2 (2.6%) |
| | Honda et al. 2020 | n = 1,366 (8.4%) | n = 6 (7.9%) |
| | Johnston et al. 2005 | n = 269 (1.7%) | n = 4 (5.3%) |
| | Macpherson and Raventos 2005 | n = 1,460 (9%) | n = 36 (47.4%) |
| | Post and Prankevicius 1987 | n = 272 (1.7%) | n = 2 (2.6%) |
| | Raventos and Macpherson 2005 | n = 402 (2.5%) | n = 4 (5.3%) |
| | Samhouri et al. 2009 | n = 515 (3.2%) | n = 8 (10.5%) |
| | Takasuka et al. 2004 | n = 131 (0.8%) | n = 1 (1.3%) |
| | Takasuka et al. 2016 | n = 1,878 (11.5%) | n = 9 (11.8%) |
| **Life event** | | | |

| | | | |
|---|---|---|---|
| | Marine migration | n = 1,1021 (67.8%) | n = 10 (13.2%) |
| | Metamorphosis | n = 2,009 (12.4%) | n = 10 (13.2%) |
| | Overwinter | n = 269 (1.7%) | n = 4 (5.3%) |
| | Post hatching | n = 316 (1.9%) | n = 2 (2.6%) |
| | Post settlement | n = 2,377 (14.6%) | n = 48 (63.2%) |
| | Young of the year | n = 272 (1.7%) | n = 2 (2.6%) |
| **Migration pattern** | | | |
| | Anadromous | n = 11,606 (71.4%) | n = 16 (21.1%) |
| | Freshwater | n = 272 (1.7%) | n = 2 (2.6%) |
| | Marine | n = 43,86 (27%) | n = 58 (76.3%) |
| **Life history stage** | | | |
| | Juvenile | n = 11,739 (72.2%) | n = 16 (21.1%) |
| | Larvae | n = 4,525 (27.8%) | n = 60 (78.9%) |
| **Species** | | | |

| | | | |
|---|---|---|---|
| | Chromis chromis | n = 816 (5%) | n = 20 (26.3%) |
| | Engraulis japonicus | n = 2,009 (12.4%) | n = 10 (13.2%) |
| | Gnatholepis thompsoni | n = 515 (3.2%) | n = 8 (10.5%) |
| | Lipophrys trigloides | n = 644 (4%) | n = 16 (21.1%) |
| | Oncorhynchus keta | n = 1,366 (8.4%) | n = 6 (7.9%) |
| | Oncorhynchus tshawytscha | n = 9,655 (59.4%) | n = 4 (5.3%) |
| | Perca flavescens | n = 272 (1.7%) | n = 2 (2.6%) |
| | Salmo salar | n = 585 (3.6%) | n = 6 (7.9%) |
| | Symphodus ocellatus | n = 198 (1.2%) | n = 2 (2.6%) |
| | Symphodus roissali | n = 204 (1.3%) | n = 2 (2.6%) |

### Figures

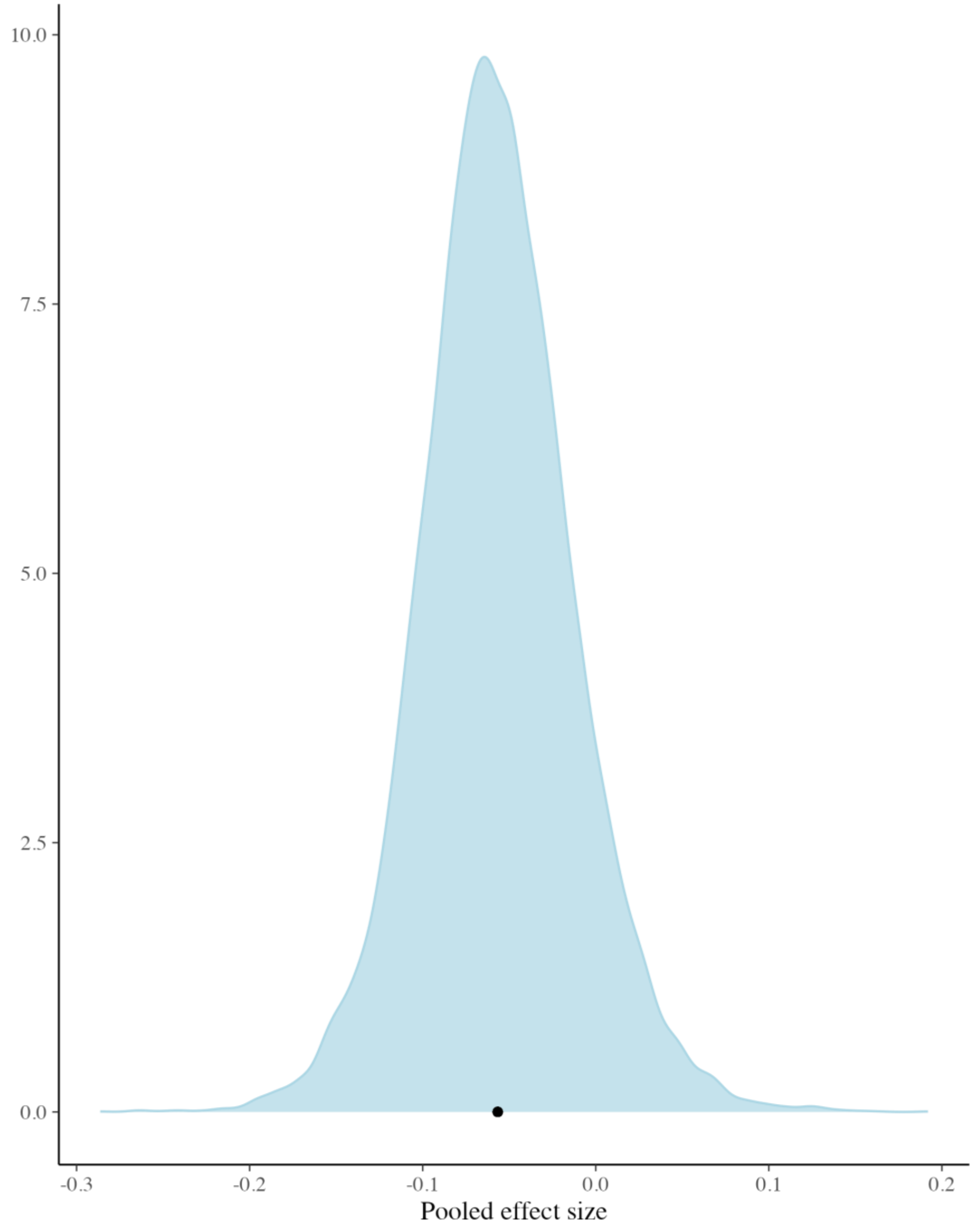

**Figure S1.** Posterior probability distribution of the pooled effect size of the original model. The black dot represents the mean of the posterior draws.

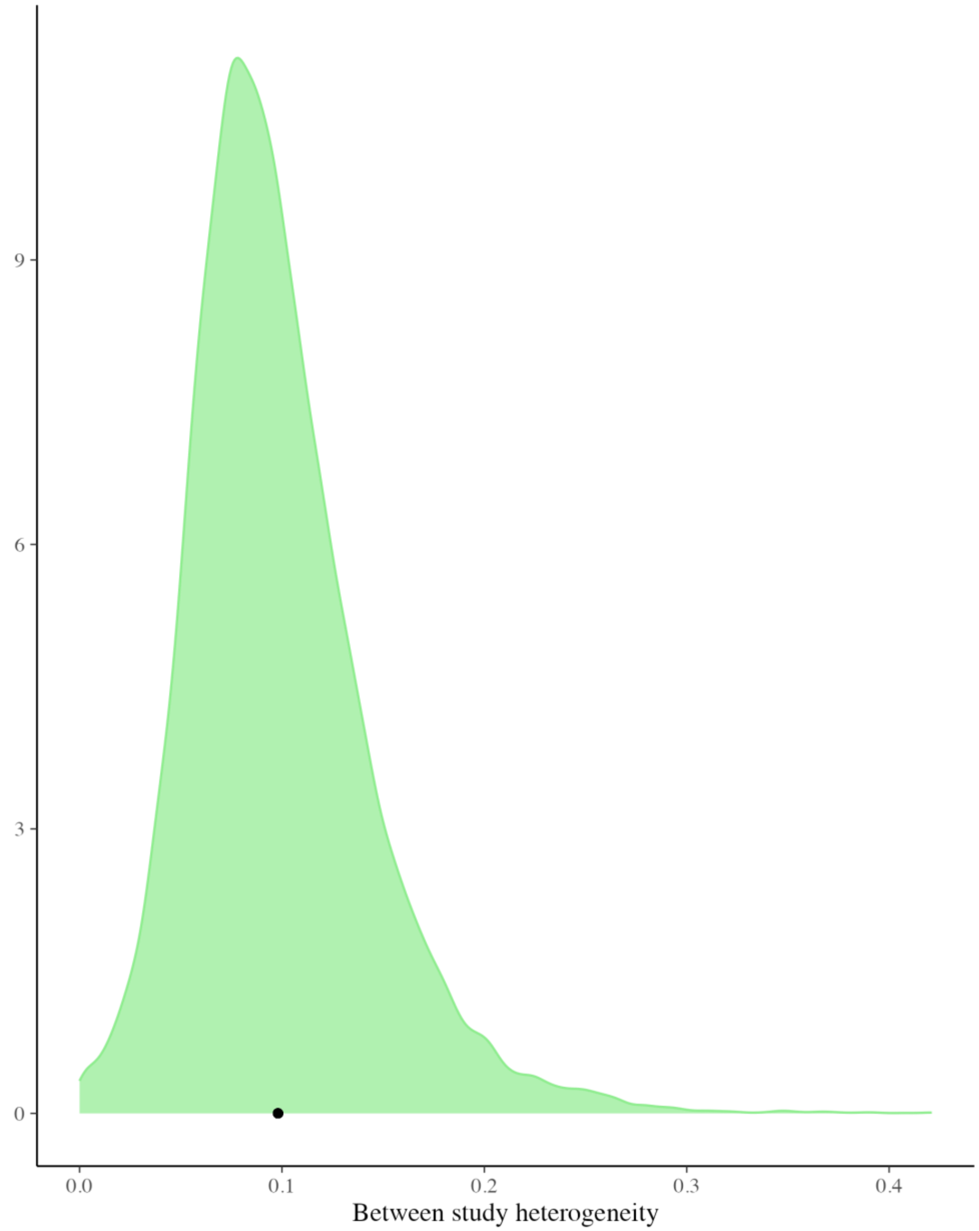

**Figure S2.** Posterior probability distribution of the between-study heterogeneity of the original model. The black dot represents the mean of the posterior draws.

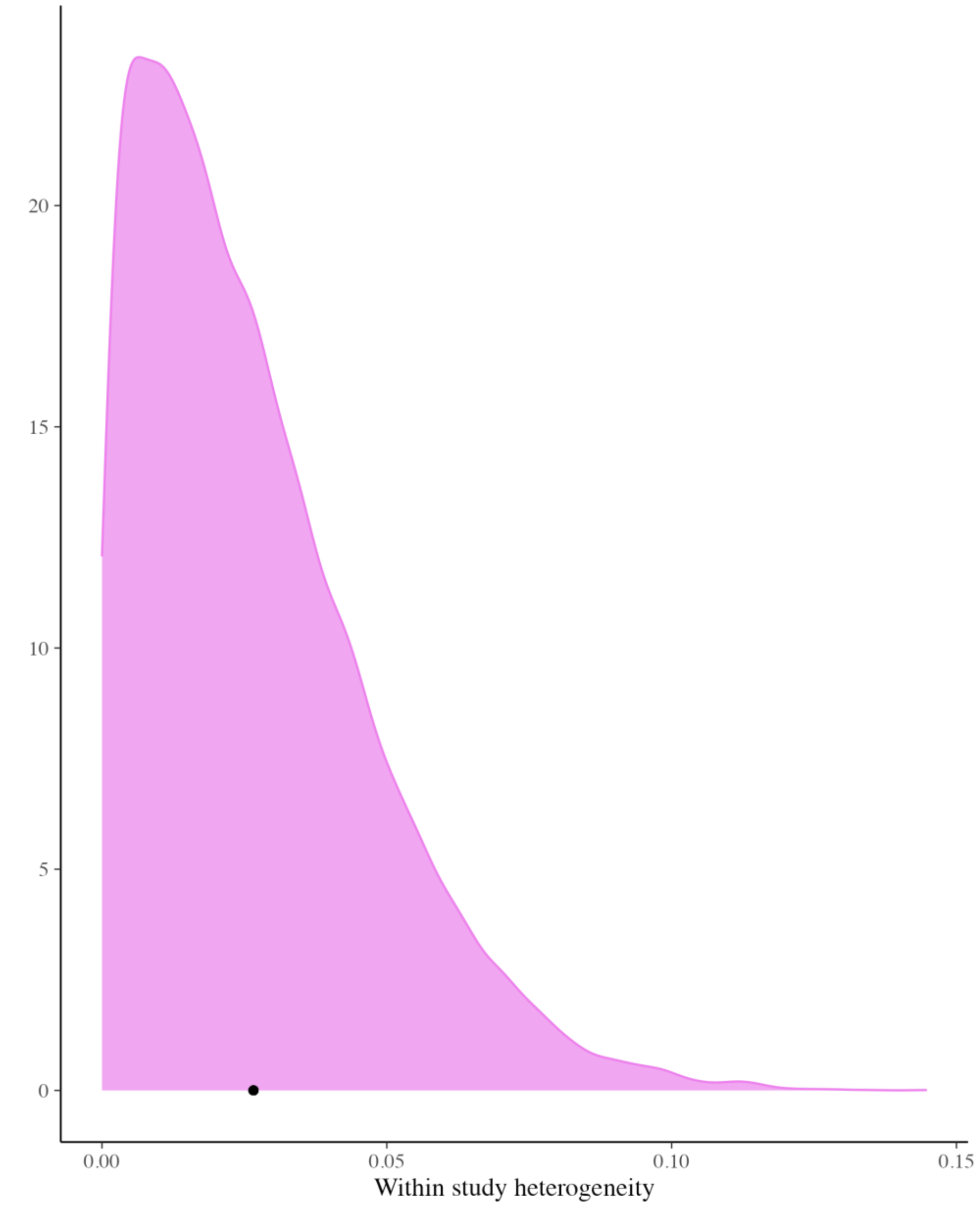

**Figure S3.** Posterior probability distribution of the within-study heterogeneity from the original model. The black dot represents the mean of the posterior draws.

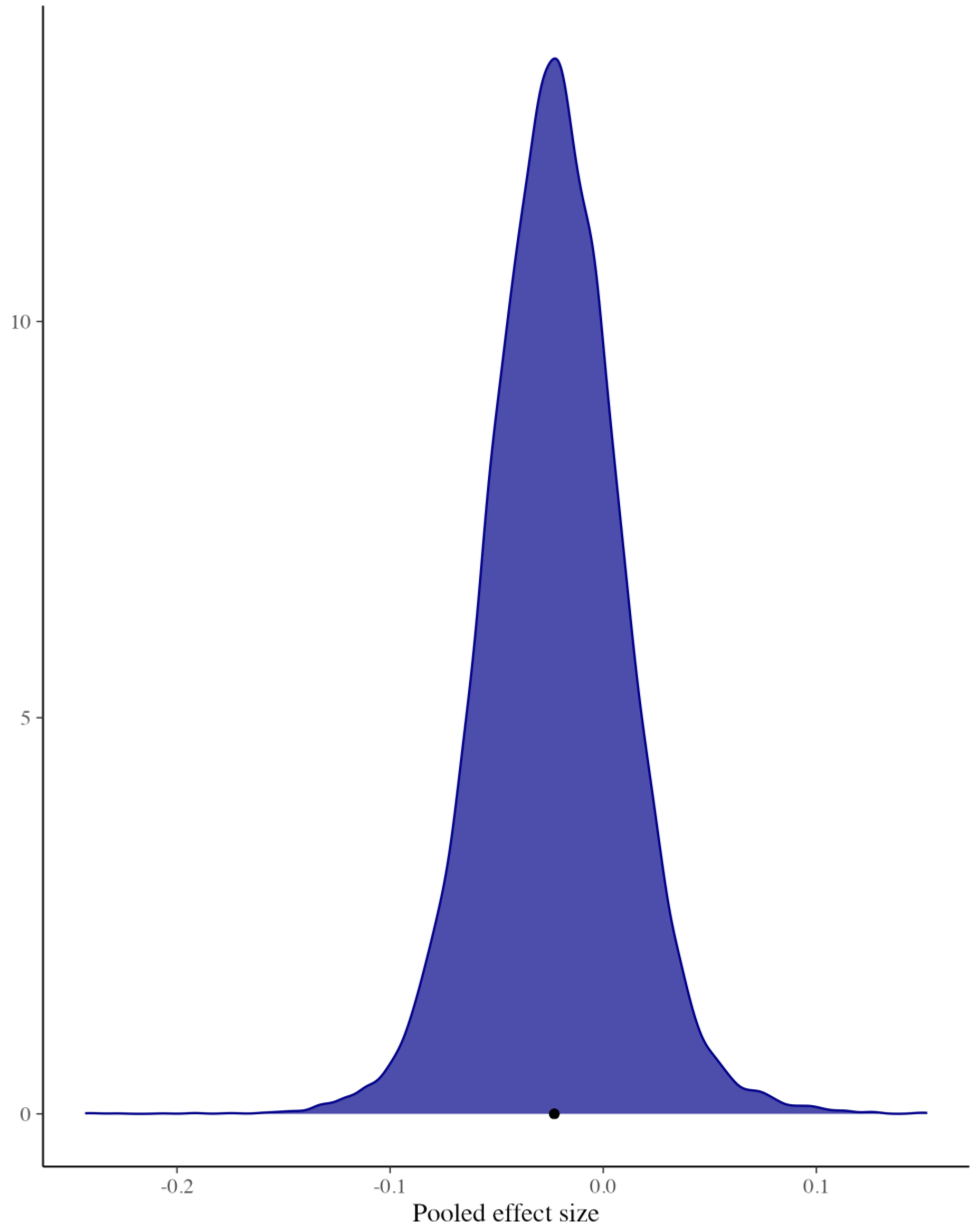

**Figure S4.** Posterior probability distribution of the pooled effect size of the sensitivity analysis model. The black dot represents the mean of the posterior draws.

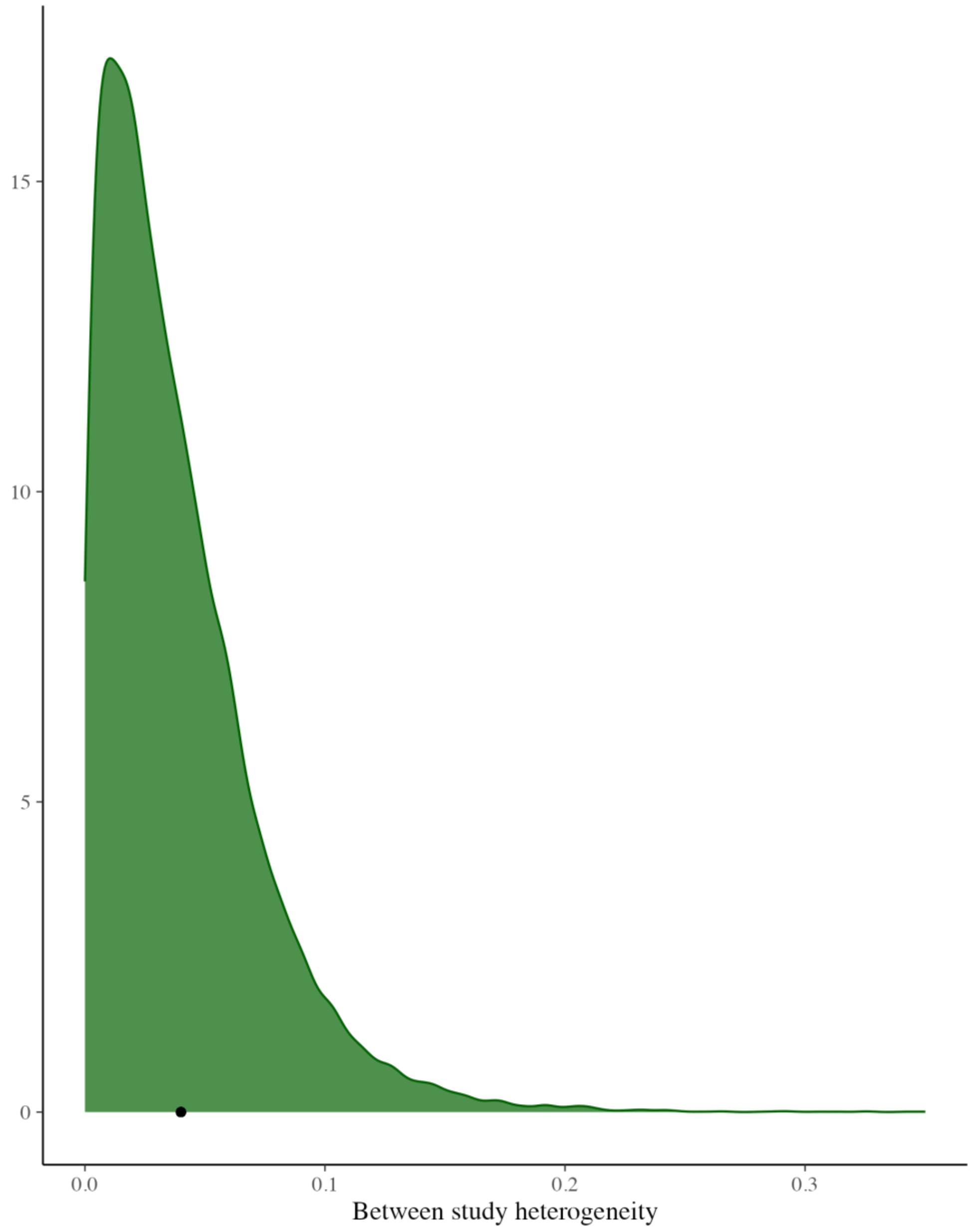

**Figure S5.** Posterior probability distribution of the between-study heterogeneity of the sensitivity analysis model. The black dot represents the mean of the posterior draws.

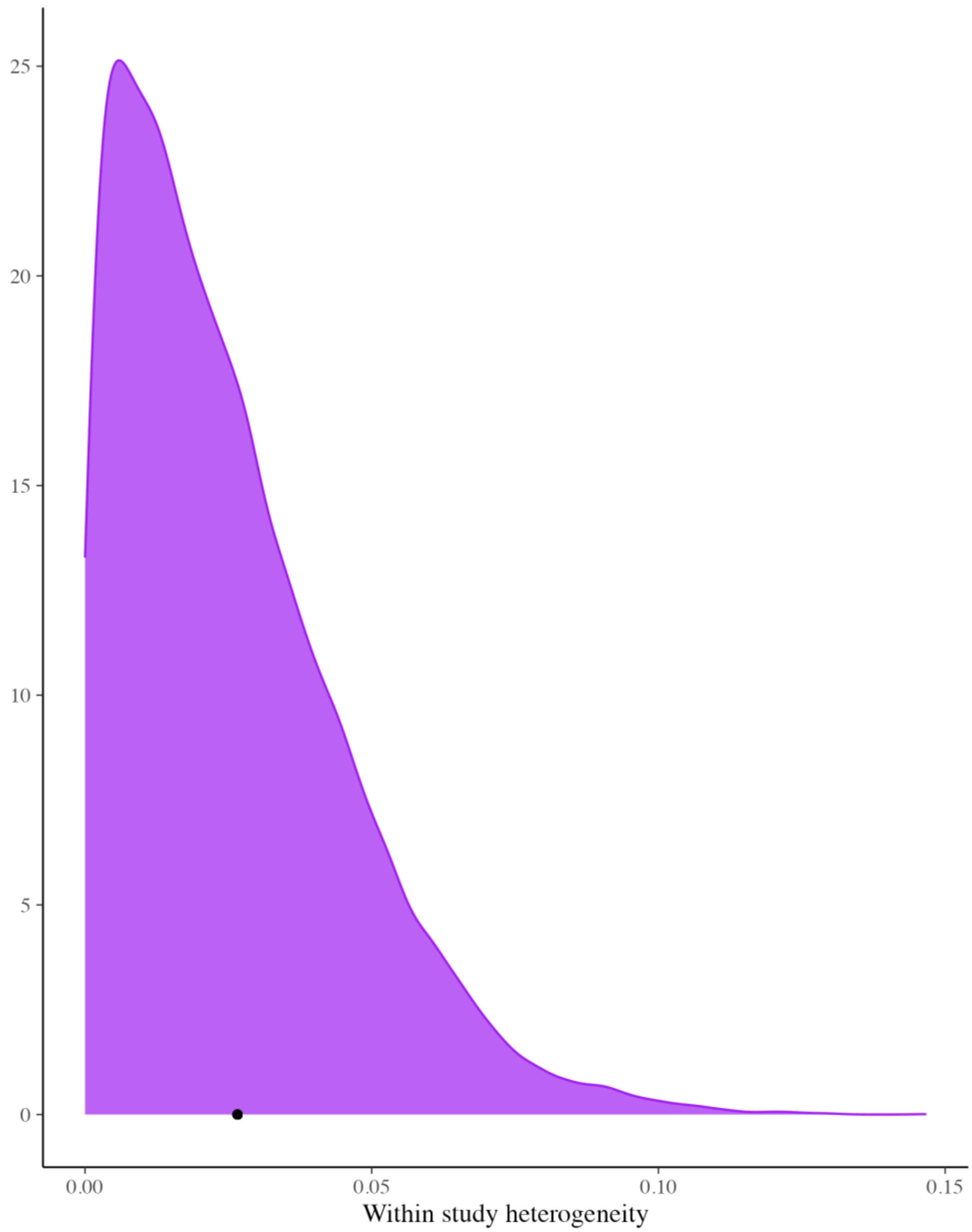

**Figure S6.** Posterior probability distribution of the within-study heterogeneity of the sensitivity analysis model. The black dot represents the mean of the posterior draws.

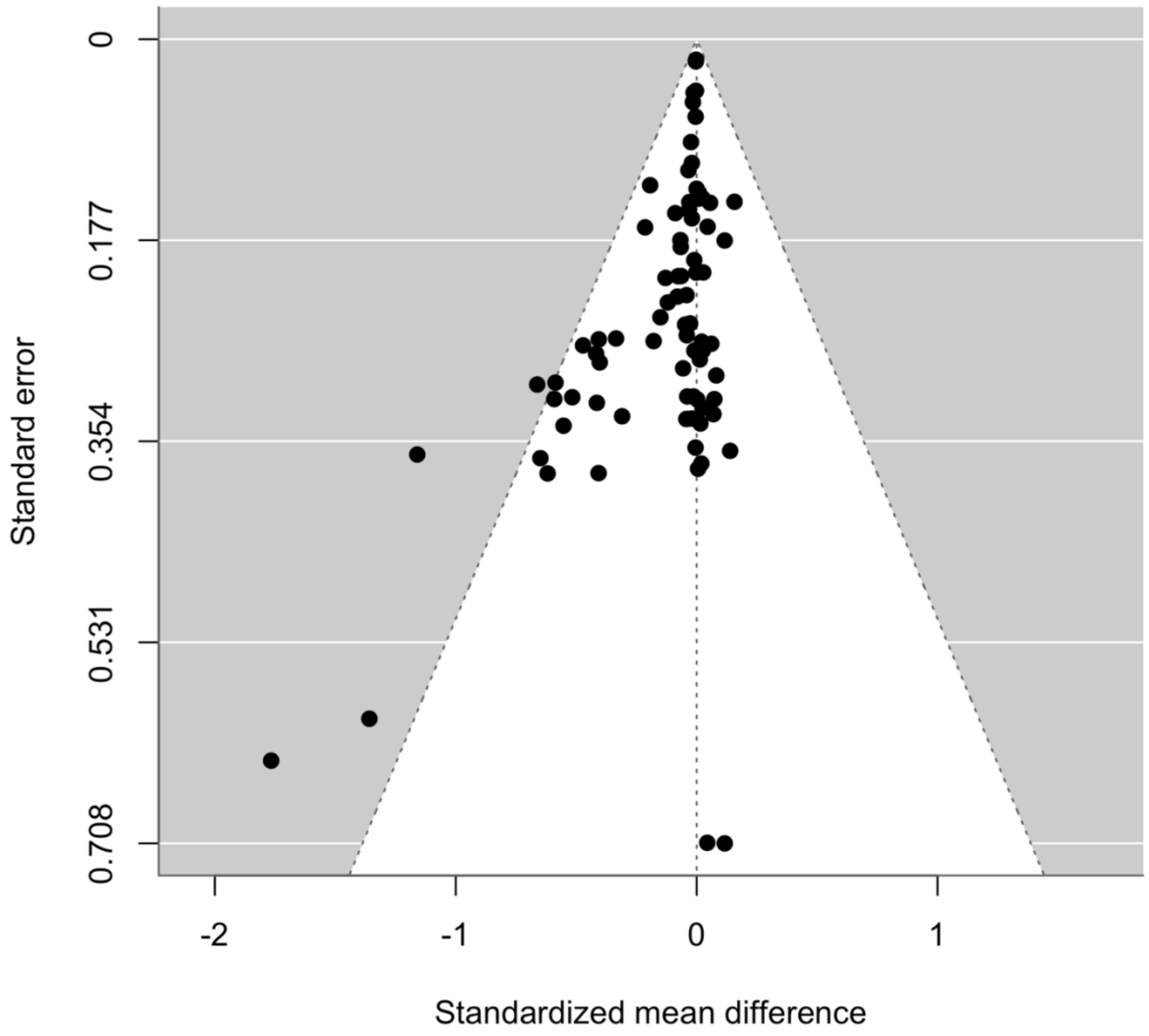

**Figure S7.** Funnel plot of 76 observed effect sizes from 10 studies showing potential bias toward publishing studies that support the 'bigger is better' hypothesis. Points represent the relationship between the standardized mean difference of the effect size and its standard error. Points outside the white funnel area indicate potential publication bias, where the standardized mean difference is larger than expected for the standard error.

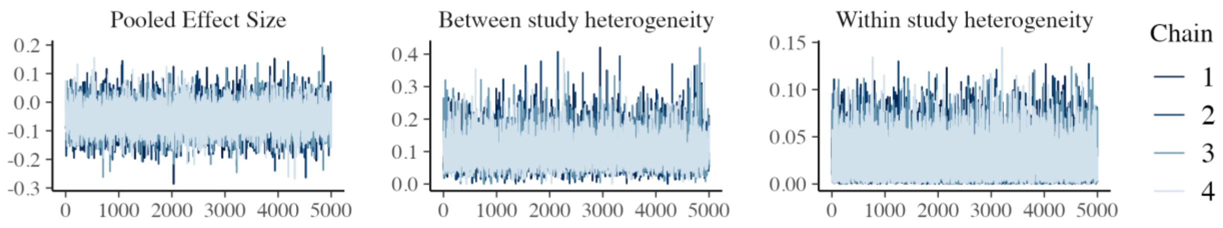

**Figure S8.** Trace plots of the pooled effect size, between-study heterogeneity, and within-study heterogeneity demonstrate adequate exploration of the parameter space by the MCMC algorithm.

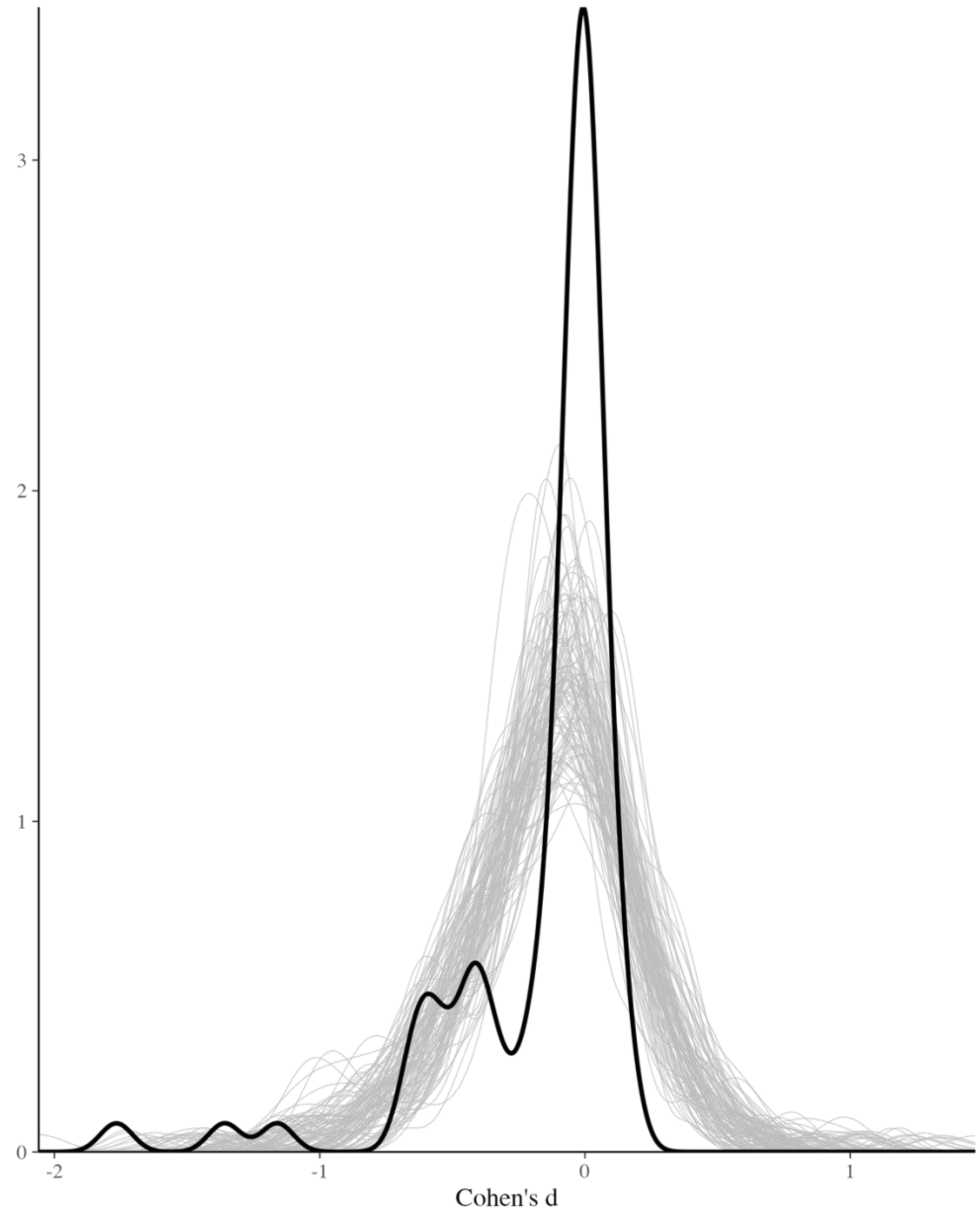

**Figure 9.** Posterior predictive check for the intercept-only model. Values of Cohen’s d simulated from the model are shown in gray, and observed Cohen’s d values are in black. Overlap between the observed and simulated data indicates a good model fit.
